# Neither sharpened nor lost: the unique role of attention in children’s neural representations

**DOI:** 10.1101/2022.08.25.505325

**Authors:** Yaelan Jung, Tess Allegra Forest, Dirk Bernhardt-Walther, Amy S. Finn

## Abstract

One critical feature of children’s cognition is their relatively immature attention. Decades of research have shown that children’s attentional abilities mature slowly over the course of development, including the ability to filter out distracting information. Despite such rich behavioral literature, little is known about how developing attentional abilities modulate neural representations in children. This information is critical to understanding exactly how attentional development shapes the way children process information. One intriguing possibility is that attention might be less likely to impact neural representations in children as compared with adults. In particular, representations of attended items may be less likely to be sharpened relative to unattended items in children as compared to adults. To investigate this possibility, we measured brain activity using fMRI while adults (21-31 years) and children (7-9 years) performed a one-back working memory task in which they were directed to attend to either motion direction or an object in a complex display where both were present. We used multivoxel pattern analysis and compared decoding accuracy of attended and unattended information. Consistent with attentional sharpening, we found higher decoding accuracy for task-relevant information (i.e., objects in the object-attended condition) than for task-irrelevant information (i.e., motion in the object-attended condition) in adults’ visual cortices. However, in children’s visual cortices, both task-relevant and task-irrelevant information were decoded equally well. What’s more, exploratory whole-brain analysis showed that the children represent task-irrelevant information more than adults in multiple regions across the brain, including the prefrontal cortex. These findings show that 1) attention does not sharpen neural representations in the child visual cortex, and further 2) that the developing brain can represent more information than the adult brain.

## Introduction

Attention is a critical system that can determine what the brain represents from the rich and complex sensory input that it receives (Posner & Petersen, 1990). Decades of research have shown that when adults attend to a particular item, the brain, especially sensory cortex, shows enhanced representations, or *sharpening*, of the attended item (e.g., more distinct neural activity patterns for the item), often at the expense of representing task-irrelevant information (Jehee et al., 2011; Kamitani & Tong, 2005). As yet, however, little is known about how such attentional sharpening develops, and how it impacts what is represented in children’s brains. Do children’s brains also sharpen representations of attended information, as has been shown in adults? Or, given behavioral evidence showing late development of attentional abilities (Enns & Cameron, 1987; Plude et al., 1994), might we observe reduced attentional sharpening and possibly greater representation of unattended information as compared with adults? Here, we address these questions using fMRI and multivariate decoding techniques. We manipulate attention by titrating the task-relevance of different stimulus dimensions —motion and objects—to measure how children’s brains represent task-relevant and task-irrelevant information.

Evidence from behavioral studies suggests that, unlike in adults, attention may not modulate neural representations in children, or at least not to the same extent. Indeed, rich behavioral literature has demonstrated that selective attention is slow to mature (Enns & Cameron, 1987; Fisher, 2019; Hanania & Smith, 2010; Plebanek & Sloutsky, 2017; Plude et al., 1994). In particular, children appear to struggle with filtering out task-irrelevant information, showing worse filtering abilities than adults in both early (e.g., 4-5yo) and middle childhood (e.g., 7-9yo), with continuous improvement until early adulthood (~18-20yos; Enns & Cameron, 1987; Hagen, 1967; Plude et al., 1994). These findings suggest that we may not observe neural evidence of attentional sharpening in children, as a possible consequence of the ongoing development of filtering abilities (Plude et al., 1994). Supporting this possibility, one neuroimaging study has shown reduced attentional modulation (i.e., lack of increasement in percent signal change through attention) in the visual cortex of middle-aged children (8-13yo) (Wendelken et al., 2011). It is therefore possible that this reduced modulation would manifest in attention having a reduced impact on how well sensory cortex represents information. However, this question of whether attention can sharpen—or even influence—the representation of attended as compared with unattended in the developing human brain is still unknown.

Alternatively, it is also possible that attention may still impact children’s neural representations, especially when children are able to pay attention to task-relevant information just as well adults. Indeed, it’s important to note that, despite ongoing development of attention, children still have significant attentional abilities: children, even young infants, can focus on particular information over others (Amso & Scerif, 2015), and young children (~4.5yo) can maintain their attention to a specific target (e.g., duck) throughout a task while ignoring distractors (e.g., turtles) (Akshoomoff, 2002). Important further work has shown that when infants expect a visual stimulus without seeing one, they display neural activation in visual cortex, just like when they are presented with a visual stimulus, consistent with the possibility that top-down signals can modulate neural activity in infants’ visual cortex (Emberson et al., 2015). Thus, attention may modulate children’s neural representations, even though such sharpening may be weaker (still improving) in children compared to adults.

These two possibilities leave questions about how the ongoing development of attention impacts neural representations in children. On the one hand, attention may have the same impact in adults’ and children’s brains, but its impact may improve as children age and become better at focusing on a target and filtering out distractors. If this is the case, when children are engaged in an attentionally demanding task and are able to perform at levels similar to adults, there should be attentional sharpening in their brains, although this could be reduced relative to adults. On the other hand, there might be a fundamental difference in the role of attention in shaping adults’ and children’s neural representations. That is, even when children can maintain their attention to particular information, there may be no attentional sharpening of relevant information, and children may represent both task-relevant and task-irrelevant information similarly. Related to this possibility, behavioral studies have shown that along with children’s poor filtering abilities, children also appear to process and represent distractor information better than adults, showing that they can remember distractors better (Plebanek & Sloutsky, 2017) as well as show better learning from distractors (Frank et al., 2021) compared to adults. That is, even when children can show sensitivity to a target, they may maintain their sensitivity to distractors, thus representing more information than adults, both relevant and irrelevant.

In the current study, we aim to explore how attention impacts neural representations in children’s brains. The behavioral evidence discussed above, which shows that children can maintain their sensitivity to unattended information, has led us to predict that unlike in adults, children’s neural representations will *not* be sharpened based on attentional relevance, even when they have adult-like sensitivity to the target. Relatedly, we also predicted that children could represent more task-irrelevant (distractor) information as compared with adults, based on behavioral evidence showing children’s greater sensitivity to distractors (Frank et al., 2021; Plebanek & Sloutsky, 2017). We explore these hypotheses using a multivariate decoding approach that characterizes distributed neural activity patterns for each type of stimulus in the visual cortex (Haxby, 2012; Haxby et al., 2001). Specifically, we scanned children ages 7-9 and adults (26 children and 24 adults included in analyses – see Methods for details) while they performed a one-back repetition detection task. All participants were asked to selectively attend to either objects (the object task) or motion direction (the motion task) and to indicate repetitions in the cued dimension. Repetitions occurred both in object and motion across all task conditions, and object and motion were always presented simultaneously and superimposed to look like a snow globe (Figure 1A & 1B). Using a support vector machine classifier, we examined how both object and motion information is represented in children’s and adults’ visual cortex when each stimulus dimension was either attended (e.g., task-relevant) or unattended (e.g., task-irrelevant). We were specifically interested in the lateral occipital complex (LOC), a brain region selective for object (Malach et al., 1995), and in the middle temporal area (MT), a brain region selective for motion information (Newsome & Pare, 1988; Tootell et al., 1995). Supporting our prediction, we found that children’s LOC and MT represent information regardless of attentional relevance, showing similar decoding accuracy for task-relevant and task-irrelevant information, whereas adults’ LOC and MT show greater decoding of object and motion, respectively, when either was task-relevant. To explore our second prediction about children possibly representing more task-irrelevant information, we examined the whole brain using a searchlight analysis to see how attended versus unattended object and motion information is represented outside of the predefined ROIs. We found that children’s brains, especially their prefrontal cortices, better represent task-irrelevant information than adults’ brains.

**Figure 1.**
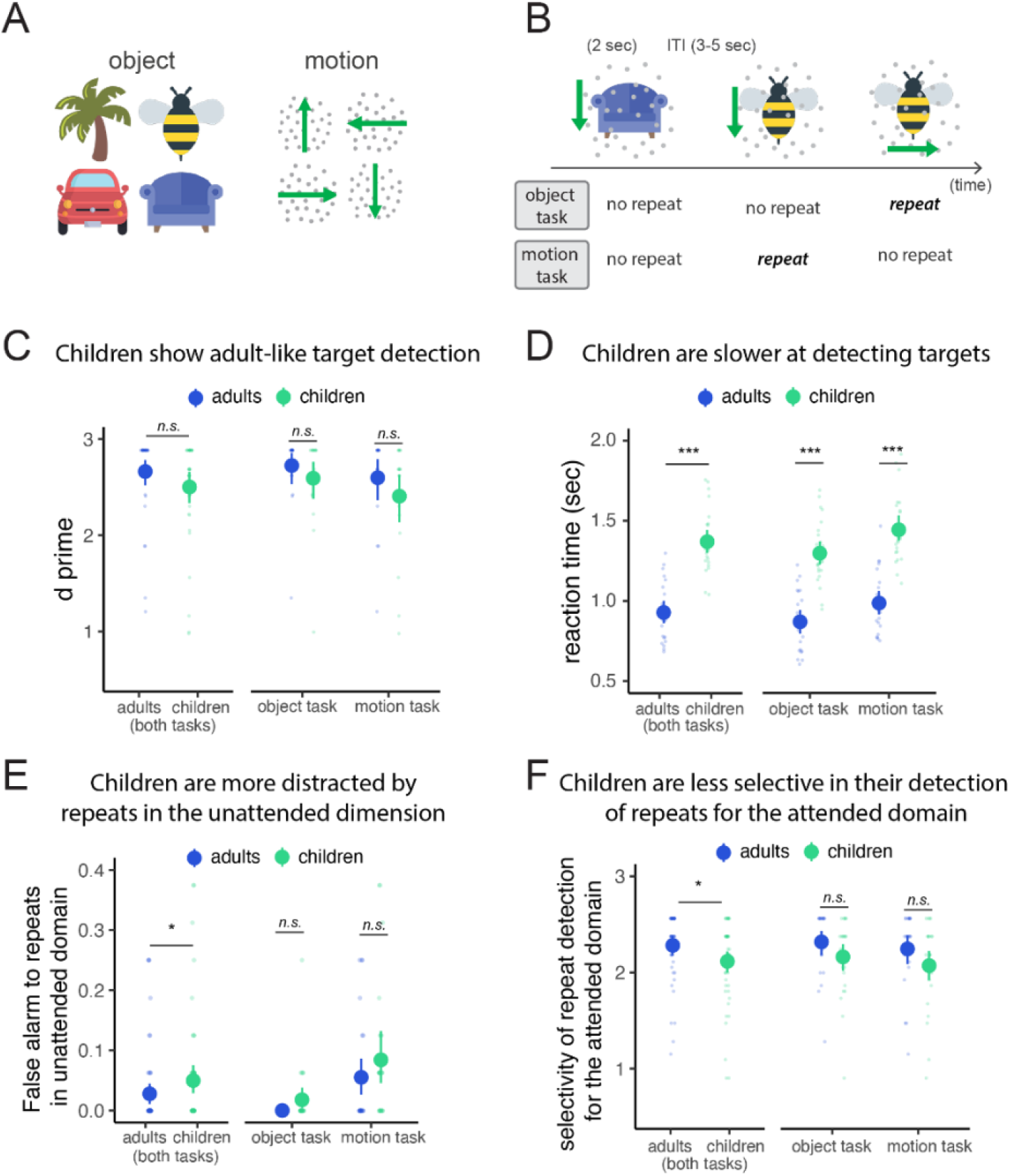
Stimuli and procedure of the experiment and behavioral performance. A) Object and motion stimuli. As shown in B), each 2-second trial consisted of one object with superimposed dots moving in one motion direction; for the object task condition (first row), participants were instructed to detect objects that repeated from the previous trial and ignore any possible repeats in motion; for the motion task (second row), participants were asked to detect repeating motion directions from the previous trial and ignore any possible object repeats. In each subsequent graph (C-F), adults’ (in blue) and children’s (in green) task performance, averaged across the task conditions, is plotted on the left, and their performance in each task condition is plotted on the right. Each colored dot indicates the mean of each group and the line indicates the estimated confidence interval (95%). Individual data are plotted in small colored dots. C) Sensitivity to target (repeats in the attended domain; d prime) for adults (in blue) and for children (in green) in object and motion task conditions, z(hit rate) – z(false alarm to all non-target trials). D) Reaction time in object and motion task conditions for adults (in blue) and children (in green). E) False alarms to repeats in the unattended domain (e.g., responding to repeats in motion in object task). F) Sensitivity to repeats in the attended domain, based on hit rate and false alarms to repeats in the unattended item (E), z(hit rate) – z(false alarm to unattended domain), *** *p* < 0.001, * p < 0.05, *n.s.* p > 0.05.

## Results

### Behavior

As shown in Figure 1C, children showed adult-like target detection performance in both task conditions. D-prime was not different in adults and children across the task conditions, *F*(1,96) = 1.294, *p* = 0.258, *np*^2^ = 0.001. Nonetheless, children showed greater sensitivity to task-irrelevant information (repeats in the unattended items) than adults (Figure 1E; left): False alarms to repeats in the unattended domain were more frequent in children as compared to adults when data were combined across the task conditions, *F*(1, 96) = 4.721, *p* = 0.032, *ηp^2^* = 0.05. Correspondingly, children showed poorer sensitivity to attended items (z(hit rate) – z(false alarm to unattended domain); see Data analyses for details) as compared to adults, *F*(1, 96) = 4.523, *p* = 0.036, *ηp^2^* = 0.04. Therefore, despite matched performance across age groups in sensitivity overall (D-prime), children’s mistakes (their false alarms) were better captured than adults’ by repeating ‘lures’ in the unattended domain, suggesting greater awareness of or processing of these irrelevant distractors. As expected (Pelegrina et al., 2015), children (mean RT: 1.32 sec, SD = 0.218) were also slower than adults (mean RT: 0.92 sec, SD = 0.187) when detecting targets across task conditions (Figure 1D; left), *F*(1,96) = 11.958, *p* < 0.001, *ηp^2^* = 0.55, both in the object, *F*(1,48) = 66.14, *p* < 0.001, *ηp^2^* = 0.58, motion, *F*(1,48) = 68.1, *p* < 0.001, *ηp^2^* = 0.59, task conditions.

### Object and motion representation in the visual cortex

Having characterized behavior, we next turned to the neural analysis. Before addressing our primary question about attention, we first aimed to establish that children’s visual cortex shows neural representation of object and motion in high specificity, showing distinct neural activity patterns for each exemplar as in adults (Kamitani & Tong, 2005, 2006), as this has never been shown before. To do so, we performed the multivoxel pattern analysis to decode object categories (bumble bee, car, chair, and tree) and motion direction (up, down, rightward, leftward) regardless whether they were attended or not, combining across the object and the motion task conditions. We used a linear support vector machine (SVM; using BrainIAK package and Scikit-learn libraries; Kumar et al., 2020; Pedregosa et al., 2011) for decoding; all runs except one run were used to train the classifier and then the remaining run was used to test the classifier (leave-one-run-out cross-validation). This analysis was performed in LOC and MT (See Methods for details on how these regions were defined). In later analyses of our primary interest, we also attempted to decode objects in MT and motion in LOC and looked at both stimuli dimensions in both regions based on attentional status.

We found sensitivity for the relevant stimulus class in both children and adults. In particular, we were able to decode objects in the LOC better than chance (25%) in both adults (mean accuracy = 41.92%; *t*(23) = 6.924, *p* < 0.001, *d* = 1.41) and children (mean accuracy = 31.97%; *t*(25) = 4.48, *p* < 0.001, *d* = 0.88; see Supplementary Figure 1A), replicating previous work in adults (Kamitani & Tong, 2005; MacEvoy & Epstein, 2009). Likewise, we were able to decode motion direction better than chance in the MT in both adults (mean accuracy = 30.45%; *t*(22) = 2.86, *p* = 0.008, *d* = 0.59) and children (mean accuracy = 32.87%; *t*(24) = 4.523, *p* < 0.001, *d* = 0.90; see Supplementary Figure 1B), again, replicating previous work in adults (Kamitani & Tong, 2006; Seymour et al., 2009).

### Attentional modulation in sensory cortex

#### Decoding in LOC

We next tested how attention modulates neural representations in the child visual cortex, and how this modulation may differ from adults, looking first at LOC and then MT. As shown in Figure 2B, in LOC, the task impacted adults’ and children’s object representations differently: there was a significant interaction between group (adults, children) and attention condition (attended vs. unattended) in the decoding of objects, *F*(1,48) = 6.983, *p* = 0.011, *np*^2^ = 0.052. Specifically, adults showed significantly greater decoding of objects when objects were attended (mean decoding accuracy when attended = 50%, SD = 18.1%; mean decoding accuracy when unattended = 33.85%, SD = 10.6%), *t*(41.83) = 3.553, *p* < 0.001, *d* = 1.025. However, in children, the attentional manipulation had no impact on object decoding, *t*(49.89) = 0.896, *p* = 0.374, *d* = 0.988 (mean decoding when attended = 34.85%, SD = 12.27%, mean decoding accuracy when unattended = 31.73%, SD = 12.86%).

**Figure 2.**
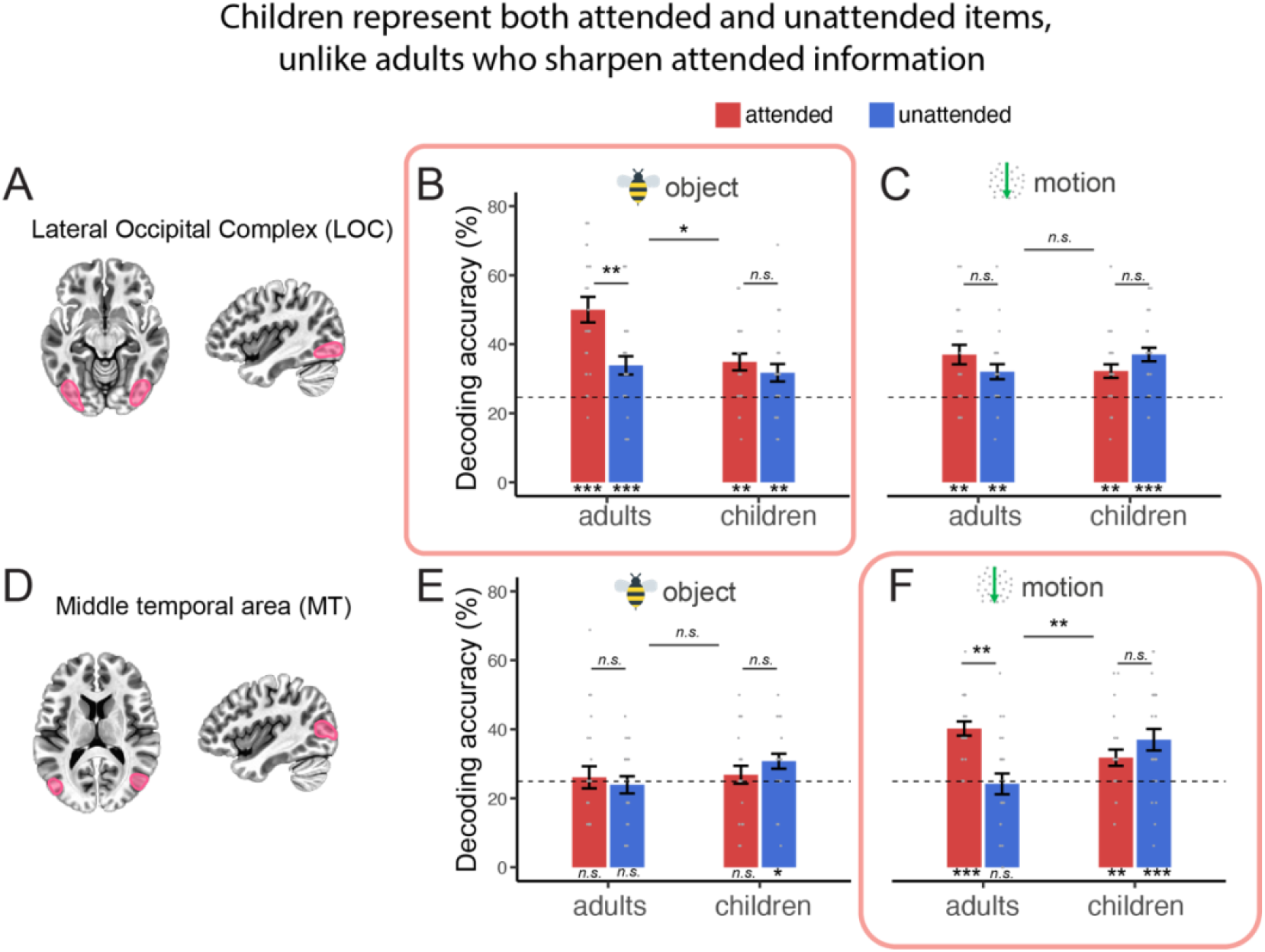
Decoding of object and motion in regions of interest. A) The Lateral Occipital Cortex is visualized, and all data plotted in this top panel (B & C) are from this region of interest. In B & C, decoding accuracy is plotted for objects (B) and motion directions (C), separately for adults and children (x-axes) when each stimulus class (objects in B, motion in C) is attended (red) or unattended (blue). Since the LOC is selective for object information, the object data are highlighted by the inclusion of a pink box around graph B. Decoding accuracy that is significantly greater than chance level is marked at the bottom of the plot (*** *p* < .001, ** *p* < .01, * *p* < .05), and significant comparisons between adults and children as well as their interactions with task condition are noted at the top of each plot (* *p* < .05). Individual data are plotted as small opaque dots, and error bar indicates SEM. For object decoding, greater decoding of attended information observed only for adults but not in children in LOC (C), and this interaction between group and attention condition is not observed for motion (C). D) The middle temporal area (MT) is visualized, and all data plotted in the bottom panel (E & F) are from this region of interest. In E & F, decoding accuracy is plotted for the four objects (E) and the four motion directions (F), separately for adults and children (x-axes) when each stimulus class was attended (red) or unattended (blue). In the MT, we observe successful decoding of motion directions only when motion was task-relevant in adults. However, children’s MT represent motion both when motion was task-relevant or irrelevant.

For motion information (Figure 2C), a stimulus class for which LOC is not specialized, there was no significant interaction between group and attention condition, *F*(1,48) = 2.534, *p* = 0.118, *np*^2^ = 0.05. Neither adults’ nor children’s decoding accuracies for motion differed by attention: adults, *t*(42.79) = 0.774, *p* = 0.442, *d* = 0.22 (mean decoding accuracy when attended = 35.15%, SD = 13.14%; mean decoding accuracy when unattended = 32.55%, SD = 9.9%); children, *t*(49.99) = −1.725, *p* = 0.091, *d* = 0.48 (mean decoding accuracy when attended = 32.21%, SD = 10.1%; mean decoding accuracy when unattended = 37.01%, SD = 9.9%). Thus, for adults, attentional sharpening of visual representation appears to occur only for objects, for which the LOC is specialized. Such attentional sharpening does not happen at all in the child LOC.

#### Decoding in MT

In MT (Figure 2D), we observed a significant interaction between group and attention condition in the decoding of motion (Figure 2F), *F*(1, 47) = 13.169, *p* < 0.001, *np*^2^ = 0.219. We found that adults showed better decoding of motion when they attended to motion, *t*(38.70) = 4.422, *p* < 0.001, *d* = 1.3 (mean decoding when attended = 40.21%, SD = 9,7%; mean decoding when unattended = 24.18%, SD = 14.38%), suggesting that attention sharpens motion representation in adult MT. Indeed, in adults, motion was not decoded above chance level when motion was not attended. In children, however, motion decoding did not differ between attention conditions, *t*(46.23) = −1.319, *p* = 0.193, *d* = −0.37 (mean decoding when attended = 31.49%, SD = 11.1%; mean decoding when unattended = 36.3%, SD = 14.89%). Thus, in adults, attention to motion sharpens its representation. This is not the case in children: attention toward motion does not impact neural representations in child MT, and child MT, unlike adult MT, represents motion direction even when motion is not relevant to the task.

For object decoding, echoing our findings of motion decoding in LOC, there was no significant interaction between group and attention condition in MT, *F*(1,47) = 2.448, *p* = 0.124, *np*^2^ = 0.05. In adult MT, object information was not represented regardless of the attention condition (attended: mean decoding accuracy = 26.08%, SD = 15.38%, *t*(22) = 0.338, *p* = 0.369, *d* = 0.07; unattended: mean decoding accuracy = 23.91%, SD = 11.86%, *t*(22) = −0.439, *p* = 0.667, *d* = −0.09). In children, however, object categories could be decoded in MT when they were unattended, mean decoding accuracy = 30.76%, SD = 10.14%, *t*(23) = 2.90, *p* = 0.003, *d* = 0.57, indicating that child MT can represent object information when adult MT does not. Interestingly, however, this object decoding in child MT was not observed when children attended to objects, mean decoding accuracy = 26.92%, SD = 12.08%, *t*(25) = 0.811, *p* = 0.212, *d* = 0.16.

Taken together, these data show that children’s visual cortex is unresponsive to attentional manipulations—decoding of the relevant stimulus class is not improved with attention. This is in stark contrast to adults’ visual cortex, which shows greater decoding with attentional sharpening. Interestingly, children also appear to represent more task-irrelevant information. Unlike adults’ MT, children’s MT represents motion information when child participants were attending to objects, which may be related to children’s greater sensitivity to task-irrelevant information (Figure 2E & 2F). In subsequent analyses, we further investigate whether children represent more task-irrelevant (unattended) information across the brain beyond these predefined ROIs in visual cortex.

### Whole brain analysis

To this end, we examined motion and object representations across the brain using a searchlight analysis. Similar to the ROI-based analysis, we here examined how objects and motion are represented when they are attended and not attended, in adults and in children. To do so, the same LORO cross-validation as in the ROI analyses was performed with the SVM classifier in small subsets of voxels (searchlights), resulting in a decoding accuracy in the central voxel of each searchlight, and it was repeated to cover all voxels across the whole brain. As we are interested in attentional relevance regardless of stimulus dimension (object or motion), we combined the data across object and motion decoding and explored how attended and unattended information are represented in adults and children. We first explored interactions between attentional relevance (attended vs unattended) and groups (adults vs children), and then we explored how adults and children represent attended and unattended information, respectively.

Echoing the findings from the ROI-based analysis where we observed greater attentional sharpening in adults, we observe several clusters showing greater attentional sharpening in adults as compared to children. Specifically, the clusters in the middle frontal gyrus (MFG), the inferior frontal gyrus (IFG), and the basal ganglia (BG) show greater attentional sharpening in adults than children (Figure 3A; Supplementary Table 1). Replicating the ROI-based findings (where we previously looked at each stimulus class separately by attention condition), we also found a cluster showing greater attentional sharpening in adults in visual cortex (Figure 3A). These findings indicate that attention does not have the same impact on children as it does on adults, not just in the visual cortex but also in the prefrontal cortex (MFG and IFG) and basal ganglia. Interestingly, children did show greater attentional modulation (e.g., greater decoding through attention) as compared to adults in the inferior parietal lobule (Figure 3A; right panel), suggesting a possible greater reliance on this earlier to develop (relative to prefrontal regions; Lenroot & Giedd, 2006) part of the association cortex for attentional sharpening processes.

**Figure 3.**
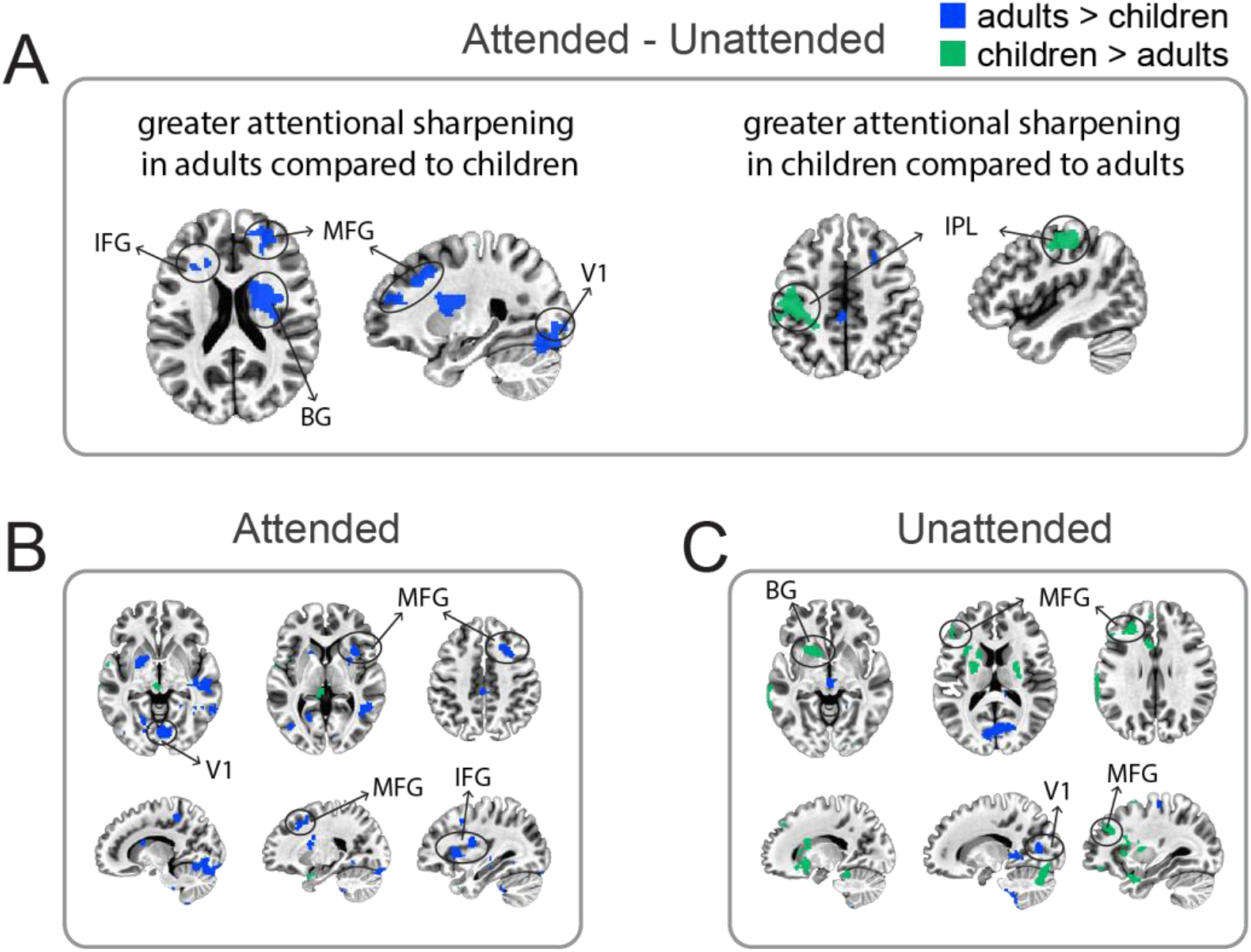
Comparison of searchlight maps between adults and children. Blue clusters indicate better decoding in adults than children, and green clusters indicate better decoding in children than adults. A) Contrast between adults and children for attentional sharpening (attended – unattended; both for object and motion). B&C) Contrast between adults and children for attended (B) and unattended (C) searchlight maps (both for object and motion). Both maps are thresholded at p < 0.05 and cluster-wise correction was conducted for multiple comparisons (minimum cluster size, 226 voxels).

To further understand how adults and children represent attended and unattended information (and not the difference between them as we described as sharpening), we compared adults’ and children’s decoding for attended and unattended information separately, again with object and motion combined.

First, for attended information, adults show better decoding than children in multiple locations across the brain, including V1, the MFG, and the inferior frontal gyrus (IFG). Indeed, most of the clusters in this contrast map show better decoding in adults than children (see Figure 3B; Supplementary Table 2). However, when information was unattended, we see a very different pattern (Figure 3C; Supplementary Table 3): Figure 3C shows quite a lot of green (greater in children) and very little blue (greater in adults). Specifically, in the anterior part of the brain, including the MFG and the basal ganglia, children show better decoding of unattended information than adults (green clusters in Figure 3C), although the opposite pattern is present in the early visual cortex (blue clusters in Figure 3C).

These findings demonstrate two interesting observations. First, although children can attend to information in a particular domain with the presence of distractors (Figure 1C), the adult brain shows better sharpening of *attended information* than the child brain (Figure 3B). Second, the child brain represents *unattended information* more than the adult brain, especially in the prefrontal cortex (Figure 3C).

## Discussion

The current study found that 7–9-year-old children’s visual cortex represents information regardless of attentional relevance, which was shown in both LOC and MT. In addition, while adults showed attentional sharpening—that is, better decoding in LOC and MT when the relevant class was attended (e.g., object for LOC)—decoding in LOC and MT was similar in children regardless of whether the relevant class was attended or not. Related to this lack of attentional sharpening in children’s visual cortex, children’s MT appears to represent task-irrelevant information, whereas this is not true for adults. Extending these findings, the exploratory whole-brain analysis shows that children’s prefrontal cortex (especially, the MFG) and basal ganglia represent task-irrelevant information more strongly than the same brain regions of adults. This greater representation of task-irrelevant information in children’s brains is likewise reflected in behavior, as children showed greater behavioral sensitivity to task-irrelevant information (Figure 2F & 2E), despite not differing from adults’ in their target-sensitivity overall. Taken together, these findings indicate that the information that is represented in children’s brains is determined by their task goals to a lesser extent than in adults; in particular, children do not prioritize or sharpen their representations of task relevant information and represent more information that is not task-relevant.

Our findings that children’s visual cortices are not modulated by attention are especially interesting on two fronts. First, these results extend previous work that shows that attention modulates neural activation in children less than adults (Wendelken et al., 2011), by showing that children are representing information very differently than adults—with reduced (or no) sharpening and greater representation of unattended information—even when children show adult-like behavioral sensitivity to targets (Figure 1C). This suggests that representing unattended information may not be costly for children, at least when the task is relatively easy. Of course, future work is needed to know more about a possible behavioral cost when the task is more demanding (say for a 2-or 3-back task). Nonetheless, these data show that even when children have adult-like sensitivity, their neural representations differ substantially in the visual cortex. As discussed in more detail below, this could have implications for understanding fundamental differences in how children process information (Gualtieri & Finn, 2022).

Second, for the first time, we address how *task-irrelevant* information in the context of attention is represented in both adults’ and children’s brains. Previous studies looking at attentional modulation in children’s brains explored attentional enhancement or sharpening for the attended information (Wendelken et al., 2011). However, no previous studies have looked at how attention impacts representations of unattended information in children. What we found using multivariate analysis is rather striking — along with lack of attentional sharpening, children’s visual cortex appears to better represent task-irrelevant information than adults’. Further, we show that beyond the visual cortex, children’s prefrontal cortex and basal ganglia represent task-irrelevant information better than those of adults. These findings together suggest children’s brains may often represent information that adults’ brains do not (Figure 2F).

These neural findings showing greater neural representation of task-irrelevant information in children are well aligned with other behavioral observations in children, which show their greater learning of and sensitivity to task-irrelevant information (Darby et al., 2021; Frank et al., 2021; Plebanek & Sloutsky, 2017; Plude et al., 1994; Sloutsky & Fisher, 2004). These findings have indicated that possibly due to the ongoing development of attention, children are more likely to process task-irrelevant information than adults, allowing them to remember or learn things that adults do not (Best et al., 2013; Plebanek & Sloutsky, 2017). While learning and memory were not measured in the present study, our findings help us understand the possible neural processes underpinning this developmental difference in learning: children’s brains indeed represent more detail about unattended information which could be called upon or used for making decisions should that information become relevant later. It should be noted, however, that much work remains to be done to link these important neural differences to differences in learning and behavior. Indeed, we did not observe a direct link between children’s sensitivity to task-irrelevant information (their greater false alarm rate to lures in the unattended dimension) and their neural representation of the irrelevant in the current study (Supplementary Figure 4). This is likely because the current study was designed to observe neural data but not optimized to observe behavioral differences in the learning of task-irrelevant information, for which a greater number of distractors and trials would be ideal. Indeed, we did not measure learning at all, something that will be critical in future work.

Along with the lack of attentional modulation in children’s visual cortex, we also show, for the first time, that children’s visual cortex represents visual stimuli with high specificity, measured with distinct neural activity patterns for different exemplars, as typically shown in adults (MacEvoy & Epstein, 2009). Previous studies looking at children’s visual cortex have mainly focused on when and how domain-selective regions emerge across development—such as the fusiform face area for faces, the parahippocampal place area for scenes, and the LOC for objects (Golarai et al., 2007; Scherf et al., 2007). These studies show that domain-specific regions in visual cortex appear to mature relatively early in life, demonstrating adult-like properties (e.g., sizes or domain specificity) in school-age children (6-8yo) (Golarai et al., 2007; Scherf et al., 2007) if not earlier (Deen et al., 2017; Kosakowski et al., 2021). Extending these previous findings, the current study shows that the LOC can also display distinct neural activity patterns for different exemplars (e.g., tree or bumble bee) within a specific domain (objects) in children, just as has been shown in adults (Haxby, 2012; MacEvoy & Epstein, 2009), and that children’s MT can represent the direction of motion, just like adults’ MT (Kamitani & Tong, 2005). Any attentional modulation effects notwithstanding, the fact that these individual exemplars can be decoded from children’s brains demonstrates that fine-grained representations of stimuli are present in LOC and MT in children, alongside their domain selectivity.

It is also important to note that our exploratory whole-brain analyses revealed insights about the development of the PFC. Along these lines, there is a large body of work which demonstrates immature PFC function through weaker neural activation related to attention, working memory, or cognitive control (Bunge et al., 2002; Crone et al., 2006; Thomason et al., 2009; Vogan et al., 2016; Wendelken et al., 2011). This paints the picture of the immature PFC as doing much less than its more mature counterpart in adults—and rightly so. Our findings add some important depth to this picture, however, by showing that the immature PFC is doing more than the mature PFC; it’s *representing* more. While this may be counter-intuitive given the especially slow maturation of the PFC (Bunge et al., 2002; Gogtay et al., 2004), these findings contribute to a growing and renewed focus on the function of the PFC in young children which shows greater functional capabilities than previously assumed (for instance, Raz & Saxe, 2020). The present work suggests that we need to rethink what maturity means from a representational perspective, in which representing less information may be more mature. Interestingly, we have known for a while that children *could* represent more than adults in immature regions, at least in theory. Indeed, a hallmark of neural immaturity is redundancy: juveniles have more redundant synapses and more neurons (e.g., thicker cortices) across the developing brain (Chechik et al., 1999; Huttenlocher, 1990), which are pruned as the brain matures. While much work is needed to establish any links in these structural terms, it could be that representing more is made possible by having more neural resources, such as more neurons or synapses.

The possible implications of the PFC representing more irrelevant information in children are far reaching. In particular, the PFC may be especially important for abstract and multi-modal representations. Indeed, recent work has shown that the adult PFC represents both acoustic and visual aspects of scenes (Jung et al., 2018; Jung & Walther, 2021) and, unlike sensory cortex, the PFC can represent stimuli more abstractly, that is independent of the modality in which they were originally presented (also see Kumar et al., 2017). This leaves open questions about what the greater representation of irrelevant information in children’s PFCs means; could the developing PFC also be representing these items in a more abstract way? And if so, how might this shape children’s learning or ability to generalize in novel circumstances? This is an interesting and important avenue for future work to better understand how representations in PFC develop.

Future investigations notwithstanding, the current study has uncovered a fundamental difference in the role of attention in shaping adults’ and children’s neural representations: unlike in adults, attention does not sharpen neural representations of attended information in children, who actually show better neural representation of irrelevant information as compared with adults. These findings are critical when thinking about how children may process and learn information differently from adults, as they reflect how information is prioritized differently in the developing human brain. Indeed, attention materialized through top-down goals and tasks appears to determine adults’ neural representations much more than children’s. Given the complexity of the world around us and the vast amount of information that we all—adults and children alike—must navigate, these findings matter a great deal. Indeed, the present data suggest that children may be more sensitive to the vast ecological complexity, and such sensitivity can be helpful and important for children when they need to learn about multiple aspects of our information-rich world at once or when their goals change. Arguably, these flexible abilities to adjust to the environment are a (or the) defining feature of childhood.

## Methods

### Participants

Twenty-six adults (mean age: 23.4 years; 281.3 months; 15 females) and 38 children (mean age: 8.9 years; 106.9 months, 18 females) participated in the current study. Of 38 children who were recruited, 10 did not complete the functional portion of the scan (mean age: 8.7 years; 104.2 months, 3 females); 7 of those children dropped out after a mock scan session, and 3 of them completed only the anatomical (T1 and diffusion spectrum imaging). Our preregistered target sample was 30 adults and 30 children (osf.io/nuf2a). And while we did not meet this goal (having 26 adults and 28 children in the final sample), we decided to halt data collection prior to meeting this goal for logistical reasons (transition of institution of the first author followed by the onset of the COVID-19 pandemic); importantly this decision was made prior to conducting analyses, both confirmatory and exploratory.

### Exclusion Criteria

Based on preregistered criteria we excluded a total of 2 adults and 2 children for: having an IQ score, measured by Kaufman Brief Intelligence Test (KBIT), that was below 80 (n = 1 adult), having missed the target on more than 60% of target trials or a false alarm rate that was greater than their hit rate (n = 1 adult who also scored below 80 on the KBIT), and having excessive motion during scanning, defined as having more than 10% of scans with higher than 2mm of framewise displacement (n=1 adult and 1 child).

Thus, 24 adults (mean age: 23.1 years/278.8 months, range: 249-373 months, 14 females) and 26 children (mean age: 8.38 years/106.4 months, range: 85-123 months, 15 females) were included in the analysis.

### Design and Stimuli

#### Attention task Runs

The experiment consisted of three different attention task conditions: the object task, the motion task, and the baseline task. For all task conditions, one of four objects (bumble bee, car, chair, tree), superimposed with dots moving in one of four directions (up, down, right, left) were present on each trial, as shown in Figure 1A. Both the object and motion tasks took the form of a one-back working memory manipulation (Owen et al., 2005), in which participants were asked to press buttons corresponding with a repeat or no-repeat on each trial. For the object task, participants were asked to find objects repeating from one trial to the next, while ignoring the motion stimuli and, of course, any repeats in the motion direction (see Fig. 1B). For the motion task, participants were asked to ignore the objects (including possible repeats) and find repeats in the motion direction (Figure 1B). Repeats did not co-occur in the object and motion dimensions; when an object was repeated, motion direction was not repeated, and vice versa. For the baseline task, participants performed an oddball detection task on the fixation cross, detecting color changes in the fixation cross. Participants were asked to press corresponding buttons (white or pink) to indicate the fixation color on each trial. Before scanning, participants first practiced the one-back task separately for objects (without motion) and motion (without object), on the same stimuli as the main experiment. Then, participants practiced the object and the motion one-back tasks, as well as the baseline task, with both object and motion presented simultaneously, just like the main experiment in the scanner. They repeated the practice until they show 75% or greater accuracy (hitting at least 3 out of 4 targets) for each task.

A mixed block/event-related design was used, where each task condition was embedded in each run as blocks, and each trial with an object image and motion stimulus was an event within each block. There were 4 total runs with three blocks per each run for each task condition and 16 trials/events per each block (exclusive pairs of four object categories and four motion directions). The order of the task blocks (object, motion, and baseline) was randomized per each run and each subject. There was a 12-second fixation block between the task blocks as well as one at the beginning and one at the end of each run.

All four object pictures subtended approximately 8.1 * 8.1 degrees of visual angle (see Fig 1A). Motion stimuli were created by using random-dot motion (RDM), which was presented within a large circular aperture (10 * 10 degrees) at the center of the screen. Each dot was approximately 0.2 degrees in diameter and moved with the speed of 4 deg/sec in one of the four cardinal directions — up, down, right, left — with 100% coherence (Figure 1A). Each dot disappeared 200 ms after its creation or when it reached the boundary of the circular area. There were always 840 dots in the display; when any dot disappeared, a new dot was created at a random location within the circular display area. Importantly, the dots and the object stimulus were always presented together, with dots overlaying the object, across all three task conditions.

#### Localizer runs

To localize object-selective regions in the brain, we performed two localizer runs where objects and scrambled-object images were presented in a block design (Malach et al., 1995). All participants performed the localizer runs after completing all four main task runs. There were 4 blocks for objects and 4 blocks of scrambled objects (18 seconds for each), and an object block and a scrambled block were always paired together, and this pair was embedded between the fixation blocks. The order of object/scrambled blocks in a pair were pseudorandomized so that for the half of the pairs, the object block came before the scrambled block and vice versa for the other half. In object or scrambled blocks, each picture of an object or a scrambled object was presented for 1 second with 600 ms of ITI. Participants were asked to watch the pictures without any explicit task. To ensure they watched all of the images, their eye gaze was monitored using an eye-tracking camera (without recording their eye movement). A 12-second fixation block was included at the beginning and end of each run.

### fMRI scanning

All scanning was performed on a 3T Siemens Prisma MRI scanner with a 32-channel head coil at the Toronto Neuroimaging Facility at the University of Toronto. High-resolution anatomical images were acquired with a MPRAGE protocol with a multiband factor of 2. Images were then reconstructed using GRAPPA, with sagittal slices covering the whole brain (T1 = 1070 ms, TR = 2500 ms; TE = 2.9 ms; flip angle = 8 deg, voxel size = 1 × 1 × 1 mm; matrix size = 256 × 256 × 176 mm). This sequence includes a volumetric navigators (vNav) prospective motion correction system, which tracks and corrects for participants’ head motion in real time (Tisdall et al., 2016). Functional images for the main and the localizer runs were recorded with a multiband acquisition sequence (TR = 2000 ms; TE = 30 ms; flip angle = 70 deg, voxel size = 2 × 2 × 2 mm; matrix size = 220 × 220 × 138 mm; multiband factor = 3; 69 slices).

### Data analysis

Prior to data collection, three adults and three children participated in a pilot version of the current study. Parameters for data analysis, including the exclusion criteria regarding head movement and spatial and temporal smoothing, were determined based on this pilot data. Data from the pilot participants are not included in the presented analysis.

#### Behavioral data

Task performance in the object task, the motion task, and the baseline task was quantified based on d-prime and the reaction time. Hits (“repeat” responses to object-repeat trials in the object task condition) and false alarms (“repeat” responses to any trials where object was not repeated in the object task condition, applying the same symmetry for the motion task condition) were recorded for all tasks. D-prime scores were calculated as *z*(hit rate) – *z*(false alarm to all non-target trials), where *z*() refers to the inverse cumulative Gaussian distribution. To better assess the sensitivity to attended items and the susceptibility to the distractors (task-irrelevant items), we also calculated the sensitivity to unattended items, based on the hits and false alarms to repeats in the task-irrelevant domain (e.g., repeats in object in the motion task condition).

We recorded reaction times (RTs) for all correct responses and excluded trials with RTs shorter than 200ms or longer than the ITI (3-5 sec, i.e., responded after the onset of next trial stimuli). Also, the RTs with larger or smaller than the mean ± 2SD within each participant were excluded. Based on these criteria, 12.97% of the trials were excluded in adults, and 22.2% of the trials were excluded in children.

#### MRI data: Preprocessing

Preprocessing of anatomical and functional data was performed using fMRIprep (version 20.0.0) and AFNI functions (version 20.3.02). Anatomical T1w images were corrected for intensity non-uniformity (INU) with N4BiasFieldCorrection distributed with ANTs 2.2.0 (Avants et al., 2014) and then skull-stripped using antsBrainExtraction (ANTs 2.2.0), followed by visual inspection for accuracy. A whole-brain mask was created for each participant using their skull-stripped anatomical T1w image for further analyses. Functional data were corrected for susceptibility distortion estimated from the fieldmap using fugue (FSL 5.0.9), co-registered to a T1w reference using bbregister (FreeSurfer), which was configured with six degrees of freedom, and corrected for head-motion using mcflirt (FSL 5.0.9). Volumes with movement > 2mm were corrected via interpolation between the nearest non-affected volumes to reduce abrupt signal changes caused by head motion (3dDespike, AFNI). No spatial smoothing was applied to the functional data of the main experiment runs. Functional data of the localizer runs were spatially smoothed with a Gaussian kernel with 4mm full width at half maximum (FWHM) using 3dmerge in AFNI. For both the main experiment and the localizer data, temporal smoothing was performed to remove frequencies above 0.2 Hz. Head-motion parameters with respect to the BOLD reference were estimated before any spatial or temporal smoothing.

#### Quality Control of Child MRI Data

We observed that children moved more than adults during the scanning. The average framewise displacement (FD) was higher in children than in adults, *t*(46.537) = 3.602, *p* < 0.001, *d* = 0.66 (Supplementary Figure 2A). However, any differences that we observe in the neural data are not likely due to differences in data quality across the two groups for the following reasons. First, in both LOC, and MT, we found that the temporal SNR (tSNR) does not differ in adults and children (Supplementary Figure 2B); in LOC, *t*(46.918) = 1.318, *p* = 0.194, d = 0.37; in MT, *t*(47.576) = 1.396, *p* = 0.1692, d = 0.4. Second, when we matched the FD values between adults and children by excluding 11 adults who stayed very still during the scanning, *t*(27.003) = 1.109, *p* = 0.276, *d* = 0.36, we still observe the same patterns of the neural data that we do without the exclusion of the adults who moved less (Supplementary Figure 2C). Finally, when we examined the univariate contrast between either of the task conditions and the baseline condition as a sanity check, we saw similar contrasts in adults and children; both adults and children show activation in the fronto-parietal regions (Supplementary Figure 3).

#### Regions of Interest

The Lateral Occipital Complex (LOC) was defined using data from the localizer runs. After preprocessing, functional data from the localizer runs were processed using a general linear model (GLM; 3dDeconvolve in AFNI) with regressors for the two types of images (object, scrambled objects) with six nuisance regressors of motion derivatives. The LOC was defined as continuous clusters of voxels with significant contrast of objects > scrambled objects, *q* < 0.05, corrected using false discovery rate (FDR) (Westfall & Young, 1993).

The middle temporal area (MT) was defined using the probabilistic atlas provided by Wang et al (2015), which was created using functional data from a large cohort of adults. First, the probabilistic MT map was thresholded at P > 10% in the MNI152 space to ensure that the atlas region is large enough to cover individual differences but still does not cross the borders between areas. The binarized map was then registered into each subject’s anatomical space using 3dNwarpApply in AFNI, which served as a template for each individual. Any voxels within the template were excluded if they were also included in the LOC mask. Within this template space, voxels were rank ordered using the GLM contrast of the functional data from the baseline condition; functional data from the baseline condition was modeled using 3dDeconvolve in AFNI with four regressors for each of four dot motion direction as well as six nuisance regressors of subject motion derivatives. We first determined the ideal number of voxels for MT by using cross-validation within the baseline condition data. Voxels were first ranked from highest to lowest F statistic of a one-way ANOVA of the activity of each voxel with dot motion direction as the main factor, and this rank order was used when selecting voxels. Then, we performed leave-one-run-out (LORO) cross validation with the number of voxels varying from 50 to 500 in increments of 50. The number of voxels that provides the best decoding accuracy was used for the main analysis (object attend condition, and motion attend condition). If there were ties, we selected the smallest number of voxels. On average, we selected XXX voxels as a result of this procedure. One adult participant did not show any motion selectivity in the MT parcel from the baseline data and thus was excluded from the MT analysis.

#### Decoding analysis

Decoding analysis was performed separately for each task condition and for each type of stimuli. First, for object decoding, we trained a linear support vector machine (SVM; using BrainIAK package and Scikit-learn libraries; Kumar et al., 2020; Pedregosa et al., 2011) to assign the correct labels to the neural activity patterns, which were the beta estimates of each object (bumblebee, car, chair, tree), inside an ROI, using all runs except one (leave-one-run-out; LORO cross-validation). The SVM decoder produced predictions for the labels of the left-out data. This cross-validation was repeated so that each run was tested once, providing predictions for object categories in each ROI and for each subject. The same procedure was performed for motion direction (up, down, right, left), resulting in decoding accuracy (a fraction of correct predictions) for object and motion in each task condition.

Group-level statistics were computed over all participants in each group (child and adult) using one-tailed t-tests, determining whether decoding accuracy was significantly greater than chance level (25% for both object and motion). The significance of the t-test was adjusted for multiple comparisons using false discovery rate (FDR). To compare the decoding performance across the condition within each group, paired two-tailed t-tests were performed. Finally, to test how decoding performance varied across conditions between adults and children, repeated-measures ANOVAs with group as a between-subjects variable, and task condition as a within-subjects variable were performed. Importantly, because we were interested in the effect of our attention manipulation (task condition) in each group, we set out to explore the simple effect of task condition (e.g., differences between attended vs unattended) in each group even when group by task condition interaction was not significant. Note that this simple effect of task condition is especially critical to better understand whether children’s brains represent task-relevant and task-irrelevant information similarly.

#### Searchlight analysis

To explore representations of objects and motion outside of the predefined ROIs, we performed a searchlight analysis using a cubic searchlight of size 7×7×7 voxels (343 voxels in volume). The searchlight was centered on each voxel within the whole brain mask, and LORO cross-validation was performed within each searchlight location using a linear SVM classifier separately for object and motion decoding in each task condition (using BrainIAK; Kumar et al., 2020). Decoding accuracy at a given searchlight location was assigned to the central voxel.

For group-level analysis, we first coregistered each participant’s anatomical brain image to the MNI 152 template using a non-linear transformation warping (3dQWarp, AFNI). We then used the same transformation parameters to register individual decoding accuracy maps to MNI space using 3dNWarpApply (AFNI), followed by spatial smoothing with a 4mm FWHM Gaussian filter. We performed one-tailed t-tests to test whether decoding accuracy at each voxel was above chance (25%) using 3dMEMA (AFNI). After thresholding at *p* < 0.05 (one-tailed) from the t-test, we conducted a cluster-level correction for multiple comparisons. We used 3dClustSim in AFNI to conduct α probability simulation for each participant. The estimated smoothness parameters computed by 3dFHWMx (AFNI) were used to conduct the cluster simulation with a p value of 0.05 as the threshold. In the simulations, a corrected α of 0.05 was used to determine the minimum cluster size. We used the average of the minimum cluster sizes (216 voxels) across all participants as the cluster threshold.

## Data and Materials Availability

The datasets generated during this study are available at https://osf.io/kd74s/

