## Supplementary materials for "Neither sharpened nor lost: the unique role of attention in children’s neural representations"

**Supplementary Figure 1**

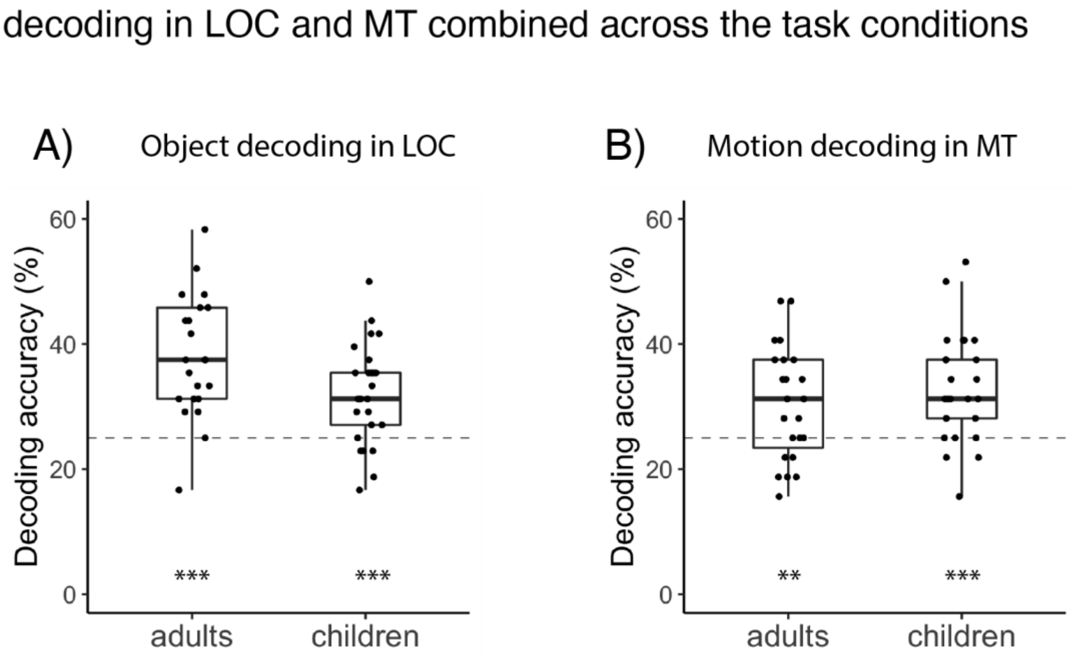


**Supplementary Figure 1** Decoding of object in LOC (A) and decoding of motion in MT (B) when data are combined across all task conditions (thus, showing decoding accuracy regardless of attentional modulation). Both adults and children show successful decoding of object (A) and motion (B). *** $p < 0.001$, ** $p < 0.01$

### Supplementary Figure 2

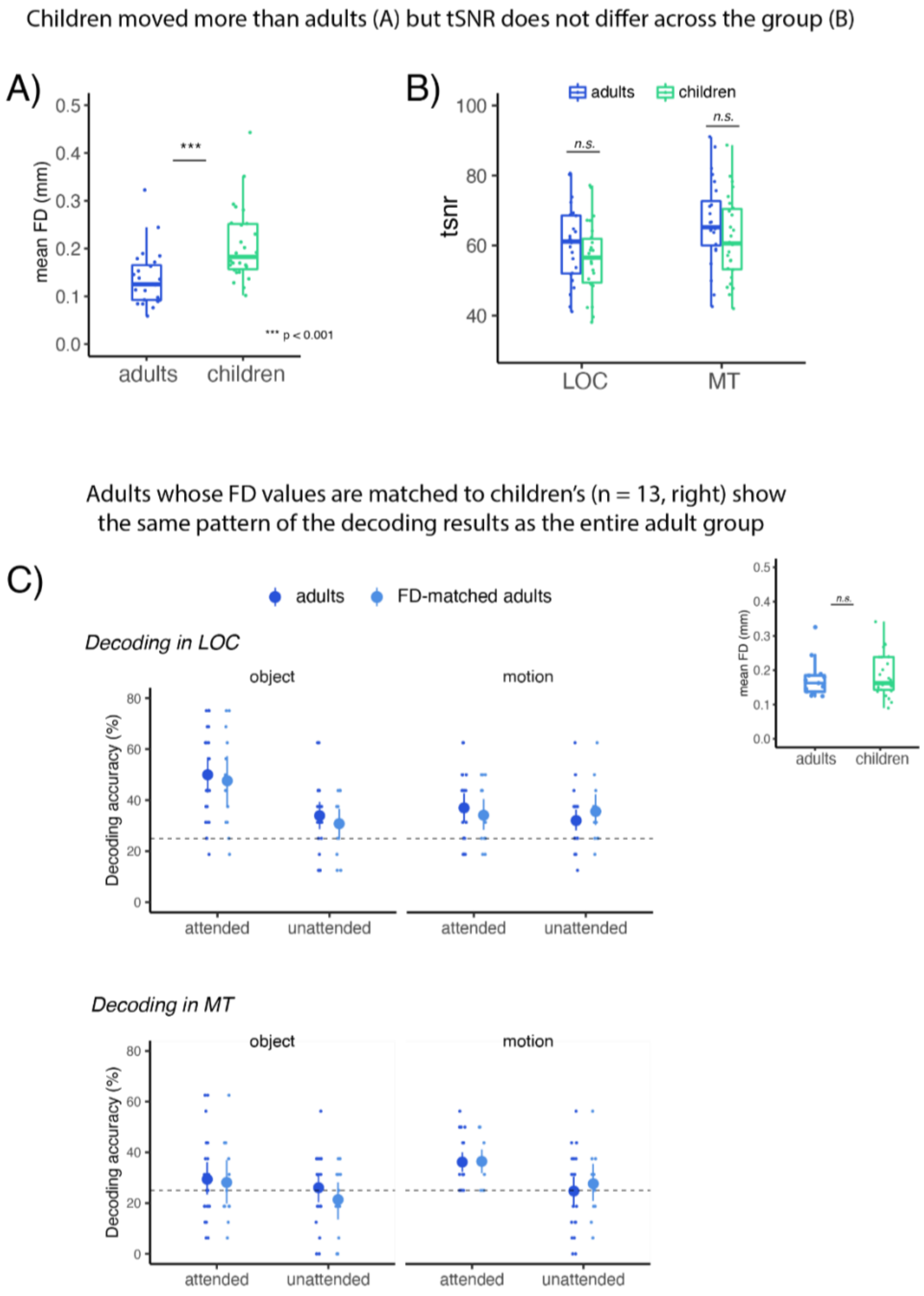


**Supplementary Figure 2** A) Average framewise displacement (FD) values in adults and children. Children showed greater FD values than adults, indicating that they moved more during the scanning. B) tSNR in the LOC and the MT. Despite greater movement, tSNR was not different in adults and children both in the LOC and the MT. A subset of adults (n=13; plotted in light blue) whose FA values are matched to those in children (plotted on the right; children's data are plotted in green) showed similar patterns of decoding as the full adult sample (plotted in darker blue).

**Supplementary Figure 3**

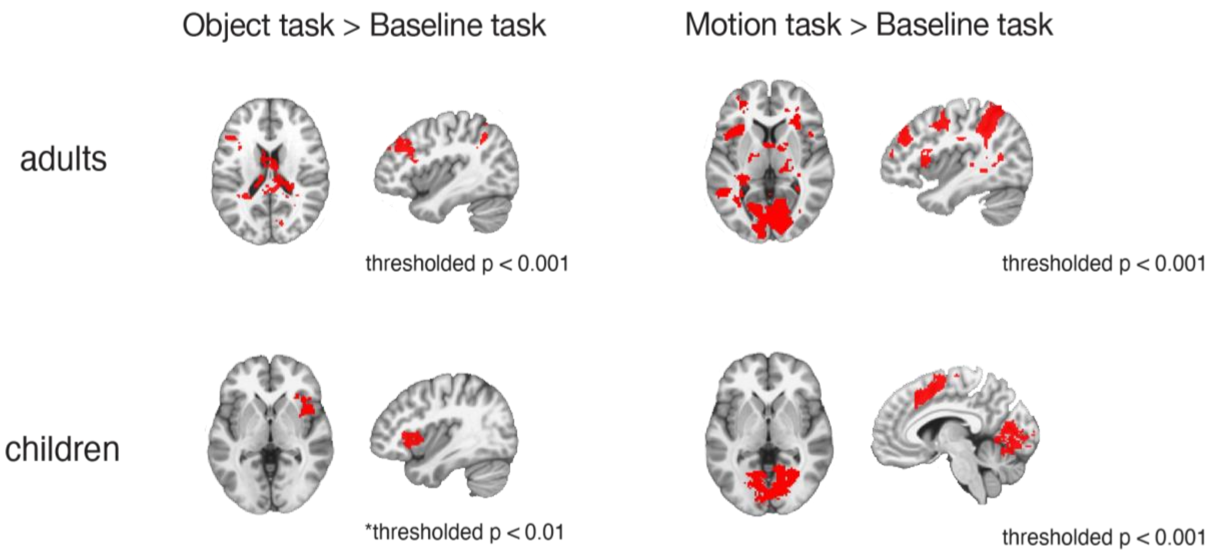


**Supplementary Figure 3** Univariate analyses showing the contrast between the object task (e.g., attending to the objects) and the baseline task (left), and the contrast between the motion task (e.g., attending to motion) and the baseline task (right), in adults (upper panel) and in children (lower panel). Adult brains show greater activation when attending to either object or motion in the fronto-parietal regions, including the middle frontal gyrus (MFG), the frontal eye field (FEF), and the superior parietal lobule, and also in the visual cortex. In children, the MFG shows greater activation in the object task. The MFG, the FEF, and the visual cortex show greater activation when attending to motion.

**Supplementary Figure 4**

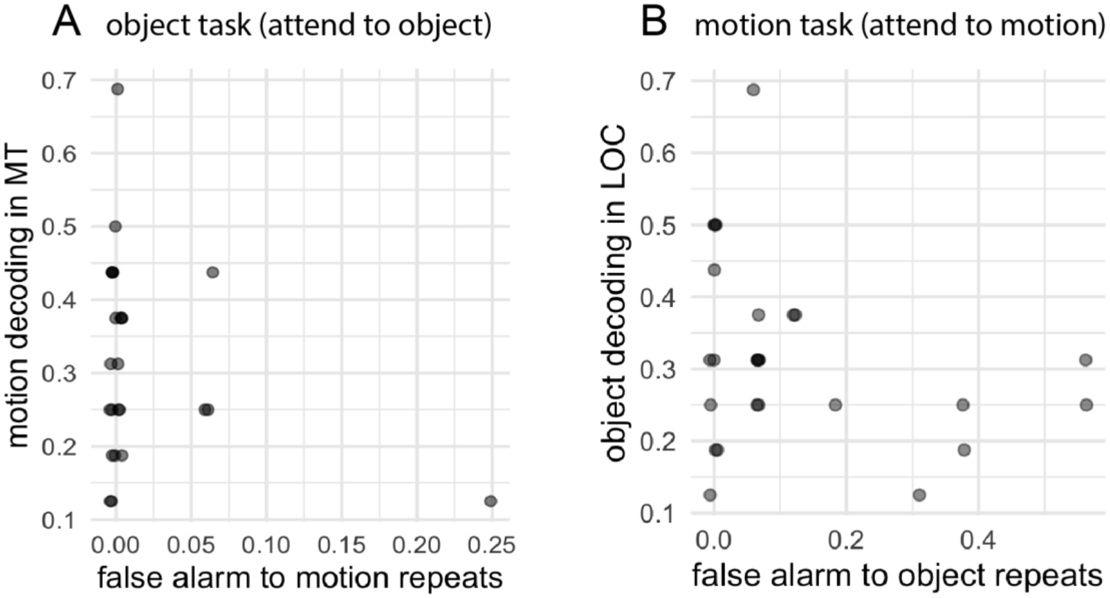


**Supplementary Figure 4** Decoding of task-irrelevant information (distractor) and false alarm to distractor in children. Individual data are plotted in opaque dots. A) In object task condition, where object was target and motion was distractor, children's false alarm to repeats in motion is plotted on the x-axis and children's neural representation of motion direction is plotted on the y-axis. Given the low number of false alarms, there is no apparent relationship ($r$ = -0.28, $p$ = 0.167) between the false alarm and decoding of motion (distractor). B) In motion task, where motion was target and object was distractor, children's false alarm to object is plotted on the x-axis and children's neural representation of objects in LOC is plotted on the y-axis. Again, there is no apparent relationship between false alarm to distractor and neural representation of distractor ($r$ = -0.282, $p$ = 0.162).

**Supplementary Table 1** Comparison between adults and children for attentional sharpening (differences in attended - unattended)

| | Peak MNI coordinate | | | Differences between adults and children (%) | Volume (ml) | Description |
|---|---|---|---|---|---|---|
| | x | y | z | | | |
| adults > children | -39.5 | 62.5 | -30.5 | 8.97 | 13648 | right middle occipital gyrus, right fusiform gyrus, right inferior occipital gyrus |
| | 20.5 | -7.5 | -10.5 | 9.05 | 9736 | left inferior frontal gyrus, left putamen |
| | -35.5 | 0.5 | 13.5 | 7.12 | 6456 | right putamen, right insula |
| | 4.5 | 34.5 | 47.5 | 6.84 | 3280 | left precuneus, left middle cingulate cortex |
| | -23.5 | -23.5 | 35.5 | 7.51 | 3216 | right cingulate gyrus, right medial frontal gyrus |
| | -31.5 | -47.5 | 17.5 | 5.69 | 2056 | right middle frontal gyrus, right superior frontal gyrus |
| children > adults | 38.5 | 30.5 | 53.5 | 7.66 | 12576 | left parietal lobule, left postcentral gyrus |
| | -7.5 | 22.5 | 65.5 | 6.84 | 2792 | right medial frontal gyrus, right superior frontal gyrus |
| | 60.5 | 4.5 | -28.5 | 5.77 | 2040 | left inferior temporal gyrus, left middle temporal gyrus |
| | 0.5 | 20.5 | -0.5 | 6.16 | 1936 | left thalamus |

**Supplementary Table 2** Comparison between adults and children for decoding of *attended information*

| | Peak MNI coordinate | | | Differences between adults and children (%) | Volume (ml) | Description |
|---|---|---|---|---|---|---|
| | x | y | z | | | |
| adults > children | -9.5 | 80.5 | -6.5 | 5.34 | 10312 | right calcarine gyrus, right lingual gyrus, V1, V2, V3 |
| | -35.5 | 2.5 | 15.5 | 3.72 | 7904 | right inferior frontal gyrus, right insula |
| | -45.5 | 52.5 | 9.5 | 4.22 | 7760 | right superior temporal gyrus, right middle temporal gyrus |
| children > adults | 52.5 | 10.5 | -38.5 | 3.64 | 3048 | left inferior temporal gyrus |
| | 32.5 | 14.5 | 69.5 | 3.85 | 2848 | left superior frontal gyrus, left middle frontal gyrus |

**Supplementary Table 3** Comparison between adults and children for decoding of unattended information

| | Peak MNI coordinate | | | Differences between adults and children (%) | Volume (ml) | Description |
|---|---|---|---|---|---|---|
| | x | y | z | | | |
| adults > children | -1.5 | 72.5 | 11.5 | 3.68 | 5536 | right/left calcarine gyrus, middle occipital gyrus |
| | 32.5 | 24.5 | 65.5 | 2.78 | 1920 | left precentral gyrus |
| children > adults | -39.5 | 64.5 | -24.5 | 3.8 | 9952 | right celebellum, right fusiform gyrus |
| | 28.5 | 4.5 | 7.5 | 3.43 | 9760 | left inferior frontal gyrus, left putamen |
| | 66.5 | 22.5 | -16.5 | 3.05 | 4328 | left middle/inferior temporal gyrus |
| | 44.5 | -19.5 | 45.5 | 3.58 | 3784 | left middle frontal gyrus |
| | 10.5 | -53.5 | -22.5 | 2.93 | 3728 | left superior frontal gyrus, left superior orbital gyrus |
| | 62.5 | 40.5 | 39.5 | 2.9 | 3640 | left parietal lobule |
